# The Genomic Architecture of Human DNA Replication Origins

**DOI:** 10.1101/2025.11.29.691293

**Authors:** Zhenhua Li, Wenjian Yang, Gang Wu, Satoshi Yoshimura, Charles G. Mullighan, Tracie C. Rosser, Elizabeth J. Leslie-Clarkson, Karen R. Rabin, Philip J. Lupo, David T. Teachey, Jun J. Yang

## Abstract

The exact sites of DNA replication origins in human and other metazoans remain elusive. Examining human whole-genome sequencing data of 2,616 specimens, we observed conjoining reads at 2,025,756 non-random genomic positions, likely arising from nascent DNA and thus defining replication origins. These origins exhibited a 16 bp motif and periodic occurrence at the intervals of 10.5 bp and 200 bp. Genome-wide replication activity is related to the expression of DNA replication-related genes. Across the genome, DNA replication initiation is more active in early replicating regions, correlated with transcription activity *in cis*, enriched for *de novo* mutations and trait-associated polymorphisms. Our high-resolution mapping of human DNA replication origins points to molecular features that govern where and when replication begins in the genome.

## Introduction

DNA replication is a fundamental biological process that ensures the accurate transmission of genetic information during cell division. To maintain genomic stability, prevent mutations, and ensure that replication occurs only once per cell cycle, the process is tightly regulated and initiates at non-randomly distributed genomic sites called “DNA replication origins” (*1*). In bacteria (e.g. *Escherichia coli*) and budding yeast (*Saccharomyces cerevisiae*), replication origins are well defined and associated with specific sequences, which are recognized by the initiator protein DnaA in the former (*2*) or origin recognition complex (ORC) in the latter (*3, 4*). In contrast, it has long been believed that metazoan genomes lack well-defined sequence-specific origins. Although tens of thousands of origins have been mapped using various genomic technologies, no consensus sequence has been identified so far. Thus, metazoan DNA replication origins are thought be influenced by a combination of chromatin context, DNA accessibility, epigenetic marks, and transcriptional activity (*5*).

The replication of eukaryotes can be divided into two steps: licensing and then firing of replication origins (*1*). In M/G1 phase, replication origins are licensed by loading the MCM2-7 helicase onto DNA with the help of CDC6 and CDT1, after recognition of origins by ORC. In S phase, licensed origins are fired when kinases such as CDKs and DDKs activate the helicase by promoting the assembly of the CMG (CDC45, MCM2-7, GINS) complex, which unwinds the DNA and enables DNA polymerase to start replication. Identification of the exact sites of DNA replication origins are complicated by multiple issues: first, only a small proportion (10% - 20%) of licensed origins are fired in each cell cycle (*6*); second, the loaded MCM complex can slide on the DNA, resulting in the actual initiation site being far away from the ORC binding site (*7*).

Previous efforts in DNA-replication origin identification, such as SNS-Seq (*8*), OK-Seq (*9*), Ini-seq (*10*), Repli-seq (*11*), have primarily relied on sequencing of newly synthesized (nascent) DNA and mapping of genomic positions by using read density-based approaches. These methods can map DNA replication origins with the size ranging from hundreds to thousands of base pairs. Such resolutions are not sufficient to pinpoint the exact site of DNA replication origins.

In this work, we hypothesize that if there are preferred initiation sites of DNA replication, they can be detected by the starting positions of nascent DNA as captured by next-generation genome sequencing. Using a whole-genome sequencing (WGS) dataset of leukemia patients (both tumor and matched germline), we identified the exact sites of DNA replication origins, identifying ∼2 million sites as potential DNA replication origins, which shared a common symmetric self-complementary motif. We further characterized the association of DNA replication origins with replication timing, genetic polymorphisms, and DNA regulatory elements.

## Results

### Identification of the exact sites of DNA replication origins in the human genome

When genomic DNA is randomly sheared for WGS, it results in sequencing reads that start and end at random positions and usually stack across a genomic region of interest. By contrast, during DNA replication, nascent DNA is synthesized from the origin bidirectionally, producing molecules that perfectly aligned at the 5’ end which leads to a series of sequencing reads with the exact start or end position. This pattern of WGS reads, therefore, is indicative of the presence of newly-synthesized DNA and replication initiation, and the positions of these reads inform the precise site of replication origins (**Fig. 1A**). To test this hypothesis, we obtained WGS data of 2,616 leukemia and germline samples from patients with T cell acute lymphoblastic leukemia (T-ALL) (*12*). WGS was performed uniformly with on ultra-sound shredded DNA fragments. For each sample, we first identified genomic positions at which a greater-than-expected number of WGS reads start or end (binomial distribution, see **Methods** for details), which were defined as significant read-starting (SRS) or significant read-ending (SRE) events, respectively. An average of 36,884 (range 805 to 1,339,857) or 36,855 (range (762 to 1,340,110) SRS and SRE events per sample were identified.

**Fig. 1.**
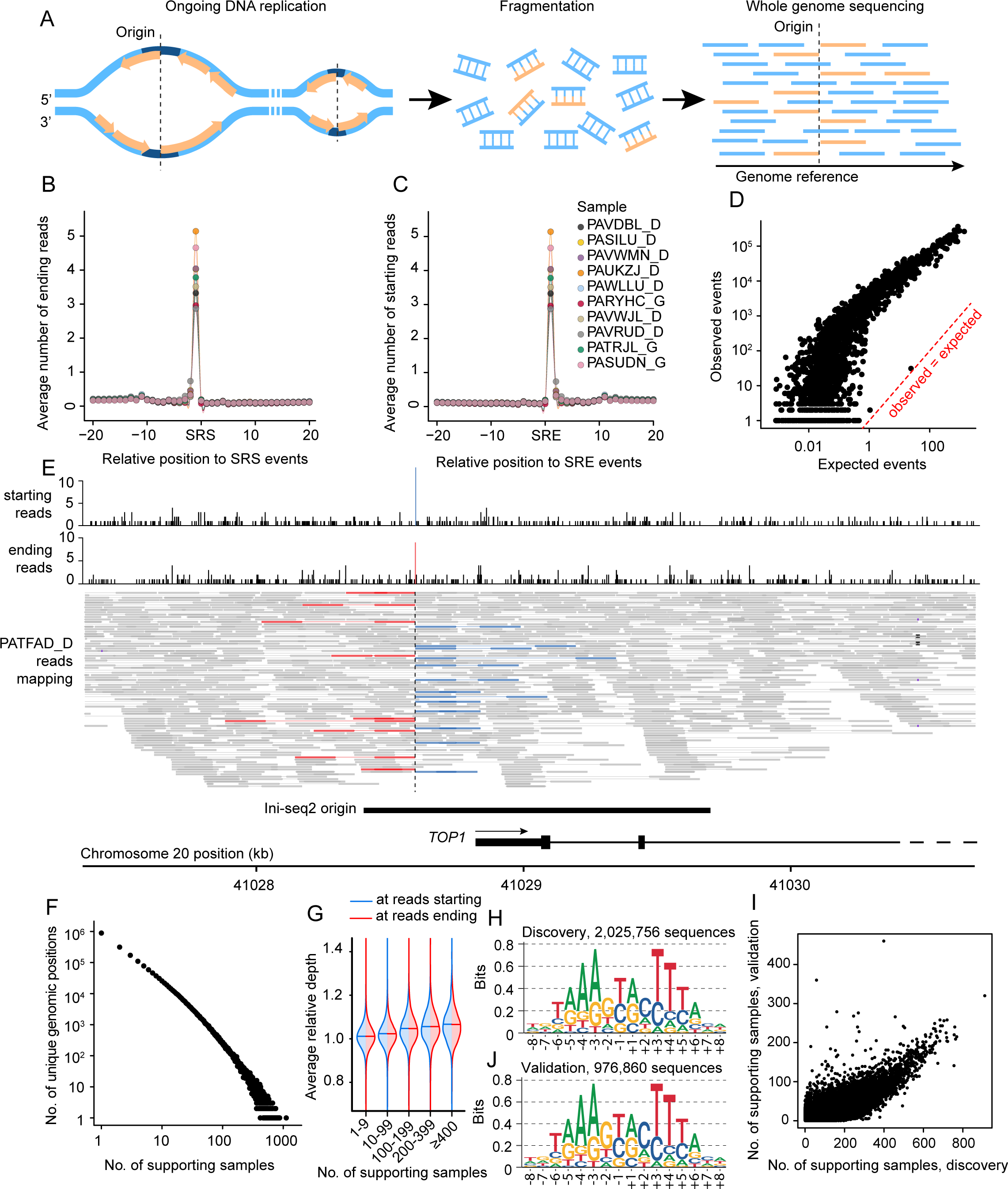
Identification of DNA replication origins from whole-genome sequencing data. (**A**) Schematic overview of the method. Unligated newly synthesized DNA results in an increased number of starting reads and ending reads at the RNA replication origin, which can be detected in genome sequencing data. (**B**) Average number of ending reads at positions relative to the SRS events. (**C**) Average number of starting reads at positions relative to the SRE events. (**D**) Comparison of the number of expected and observed SRE-SRS pairs. (**E**) Example of the detection of an SRE-SRS event in a sample. (**F**) Distribution of the number of samples in which an origin was detected. (**G**) Distribution of the average relative depth of the origins grouped by the number of samples in which they were detected. (**H**) The motif of the sequences around the DNA replication origins in the discovery dataset. (**I**) Comparison of the number of samples the origins are detected in on the discovery and validation data. (**J**) The motif of the sequences around the DNA-replication origins in the validation dataset. Created in BioRender. Li, Z. (2025) https://BioRender.com/36bnyss (**A**).

An exceedingly high number of reads ended immediately upstream of SRS events, and an exceedingly high number of reads started immediately downstream of SRE events (**Fig. 1B** and **1C**), and this unique paired pattern suggests that these events originate from the same positions in the genome. We thus posited that an SRE-SRS pair (with SRS immediately following SRE) define the exact sites of DNA replication origin. Cumulatively in 2,612 samples, 13,843,609 SRE-SRS pairs (median = 7; range 0 to 362,603) were detected, which are mapped to 2,025,756 unique genomic positions, averaging 1.56 origins per kb. We estimated the expected number of origins detected under the assumption that SRS and SRE events were independently distributed in the genome. The median observed-to-expected ratio was 293, implying bidirectional progression of replication forks from replication origins as evidenced by the pairwise occurrence of SRS and SRE events (**Fig. 1D**). More replication origins were identified in tumor samples than normal samples, likely due to the higher homogeneity of tumor cells (**Fig. S1**). An example is depicted in **Fig. 1E**, where a replication origin was identified in the well-known DNA replication origin located near the 5’ UTR of *TOP1* gene (*13*).

Among the analyzed samples, the number of times a replication origin was detected followed the power law (**Fig. 1F**). The majority of the sites were detected only in a few samples, consistent with the stochastic nature of DNA replication origin firing. 43.9% of the origins were detected in only one sample, and 0.8% of the origins were detected in more than 100 samples.

To show that our inferred replication origins were not an artifact of biased DNA fragmentation during WGS library construction, we examined the average depth of coverage at the origins relative to positions randomly sampled across the genome. Increased coverage was seen in identified DNA replication origins, with an average increase of coverage from 1% for those origins appearing in <10 samples to 7% for those appearing in >400 samples (**Fig. 1G**).

Examining the sequence at the identified replication origins, we observed a highly-conserved 16 bp symmetric palindromic motif (**Fig. 1H**). This is consistent with the fact that DNA replication origin firing is bidirectional. From the origin, we numbered these positions as forward 1 to 8 (+1 to +8) and reverse 1 to 8 (−1 to −8) based on the sequence orientation of the hg38 genome reference. Strong preferences of base usage were observed at these positions. Specifically, A or T was rarely seen at positions +1 or −1, respectively; while G or C was rarely used at positions +3 or −3, respectively. The absence of nucleotide bases at these positions and the symmetric nature of the motif also indicates that these sequences are not the junctions of Okazaki fragments which are synthesized in a uni-directional fashion.

To validate these findings, we obtained an independent WGS dataset from individuals enrolled in the Gabriella Miller Kids First Pediatric Research Program Down Syndrome (DS), Heart Defects, and Acute Lymphoblastic Leukemia study. This dataset consists of a total of 1182 samples, including normal and tumor samples from DS individuals with B-cell acute lymphoblastic leukemia (B-ALL), as well as samples from DS individuals with congenital heart disease and their parents (see **Methods**). Similarly in this validation dataset, the number of observed SRE-SRS pairs were significantly higher than expected (**Fig. S2A**). Overall, in the validation dataset, 2,848,989 SRE-SRS pairs were identified, defining 976,860 unique origins. The prevalence of each replication origin across samples followed a similar pattern of distribution as in the discovery cohort (**Fig. S2B**), with the number of detected shared origins decreasing exponentially with the number of samples. Generally, the origins detected in more samples in the discovery set were also detected in more samples in the validation set (**Fig. 1I**). This suggests that though the firing of origins is stochastic in general, some origins are more frequently used than others. The motif of the replication origins was also consistent between the validation and discovery cohorts (**Fig. 1J**).

In the following sections, we investigate the properties of the origins, focusing on the sites identified in the primary discovery dataset.

### Periodic occurrence and uneven distribution of DNA replication origins in the genome

To understand the distribution of DNA replication origins in the genome, we examined the distance between adjacent origins across the genome (**Fig. 2A**). Overall, replication origins were frequently positioned with 200bp intervals. Within the 200bp-window, hotspots for replication origin were seen roughly 10.5bp apart from each other. We formally estimated the probability of observing another origin from a given genomic position of origin and confirmed this periodic pattern (**Fig. 2B**). These observations suggest that the occurrence of DNA replication origins in the genome is periodic, with a major interval of 200bp, and a minor interval of ∼10.5bp.

**Fig. 2.**
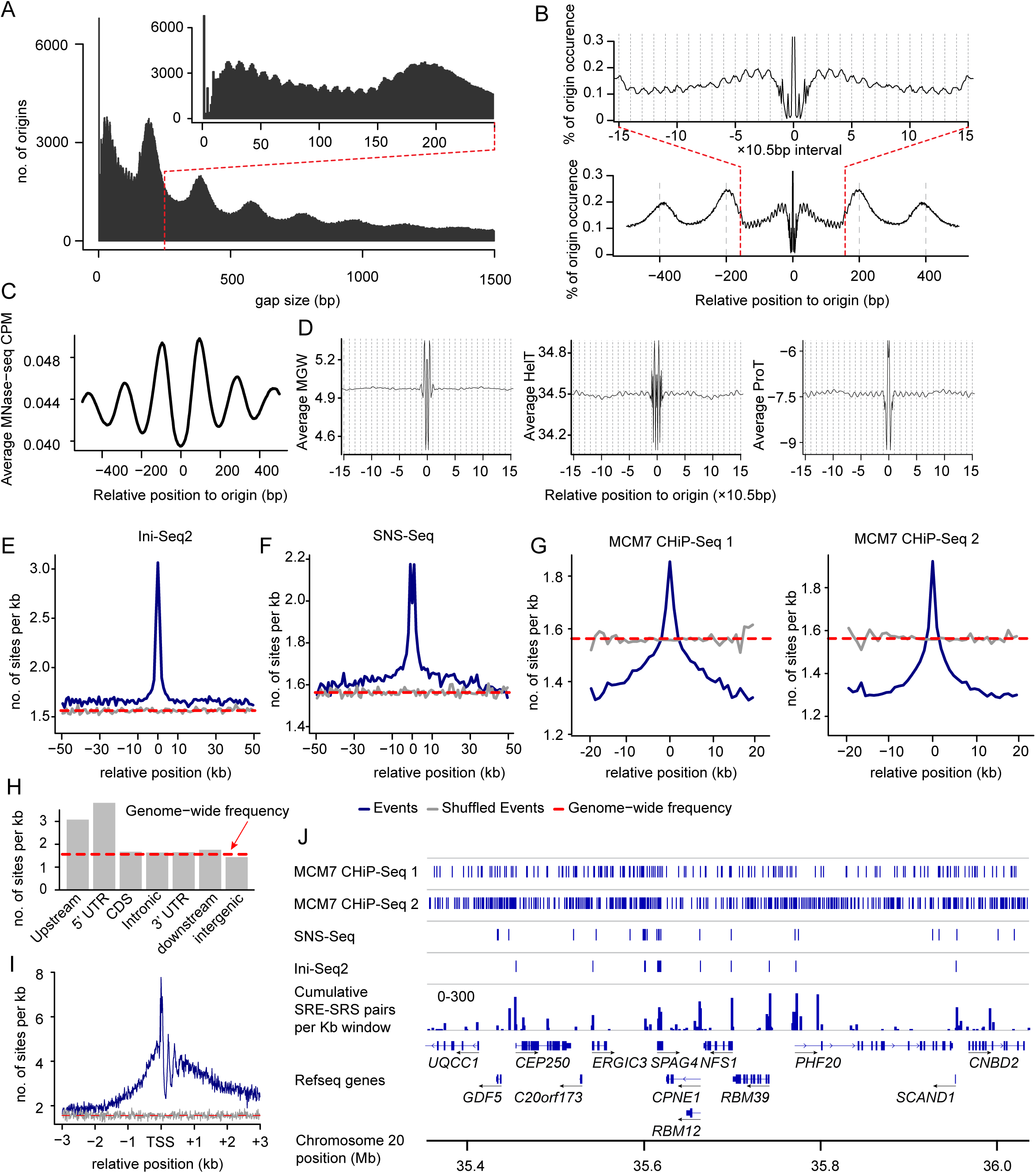
Distribution of DNA replication origins in human genome. (**A**) Histogram of the gap between adjacent origins. The inset shows the distribution within 300 bp. (**B**) Probability of detecting another origin around the origins. (**C**) Average signal (Counts Per Million, CPM) of MNase-seq at varied positions relative to replication origins. (**D**) Distribution of the average minor groove width (MGW), helix twist (HelT), and propeller twist (ProT) in regions adjacent to replication origins. (**E)-(G)** Frequencies of DNA replication origins vary by distance to regions identified by Ini-Seq2 (**E**), SNS-Seq (**F**) and MCM7 CHiP-Seq (**G**). (**H)** Frequencies of DNA replication origins in different types of genomic regions relative to genes. (**I**) Variations of the frequencies of the DNA replication origins relative to the distance to transcription start sites (TSS). (**J**) Illustration of the distribution of identified DNA replication origins, Ini-Seq2, SNS-Seq, and MCM7 CHiP-Seq in a region on chromosome 20.

Because the major interval is roughly the size of a nucleosome-linker repeat, we hypothesized that DNA replication preferentially starts at the linker between the nucleosomes due to greater chromatin accessibility. To confirm this, we obtained previously published MNase-Seq data (*14*), and calculated the average signal strength of nucleosome binding around our identified origins (**Fig. 2C**). The nucleosome binding signals aligned with the replication origins, with the lowest nucleosome binding observed at the origin, and at positions with distance in multiples of 200bp, suggesting that DNA replication origins are most likely to be within the linkers between adjacent nucleosomes.

The length of the minor interval (∼10.5bp) is approximately the size of one complete turn of B-form DNA, which suggests the local structural requirement of the DNA replication origin. To investigate if there is a recurring pattern of DNA structure around the origin, we used the computationally annotated DNA shape database GBShape (*15*), and obtained minor groove width (MGW), helix twist (HelT) and propeller twist (ProT) of the positions within 140bp of the origins. Large fluctuations of these indices are seen within the ±8bp region, likely related to the local DNA structure within the replication origin motif. Moreover, these indexes, especially HelT and ProT, displayed periodic change per 10.5bp around DNA replication origins (**Fig. 2D**).

Further, we compared our inferred replication origins with genomic regions previously identified by Ini-Seq2 (**Fig. 2E**) and SNS-Seq (**Fig. 2F**), as well as MCM7 CHiP-Seq (**Fig. 2G**) (*8, 10, 16*). Notably, DNA replication origins identified in this study were much more likely to be located within or proximal (<5 kb) to previously reported regions using these methods.

Across genes, upstream (within 2 kb) and 5’ UTR regions exhibited increased frequencies of DNA replication origins (**Fig. 2H**), with a marked enrichment around the transcription start site (TSS) of protein coding genes (**Fig. 2I**). In the upstream region of TSS, frequencies of DNA replication origins decreased gradually as the distance increased, while in the downstream of TSS, a periodic pattern of DNA replication origin frequency was observed, likely due to the phased occupancy of nucleosomes in 5’ UTR (*17*) (**Fig. 2I**). An example of this pattern on Chr 20 is shown in **Fig. 2J**.

Taken together, these data suggest that although the firing of replication origin is stochastic, there is considerable conservation of replication origin usage across different cell types.

### DNA replication activity is associated with the expression of replication-related genes

We posit that replication activity at each origin is reflected by the number of WGS reads that start at the +1 position or end at the −1 position relative to the replication origin (**Fig. 3A** and **Methods**), and the genome-wide abundance of these reads indicate the overall DNA replication activity for that individual. We observed a near-perfect correlation between replication activity estimated from reads starting at +1 vs ending at the −1 position (**Fig. S3A**), consistent with the fact the firing of replication origins is simultaneously bidirectional. Due to this high correlation, we focused on starting reads-based activity for the following analysis.

**Fig. 3.**
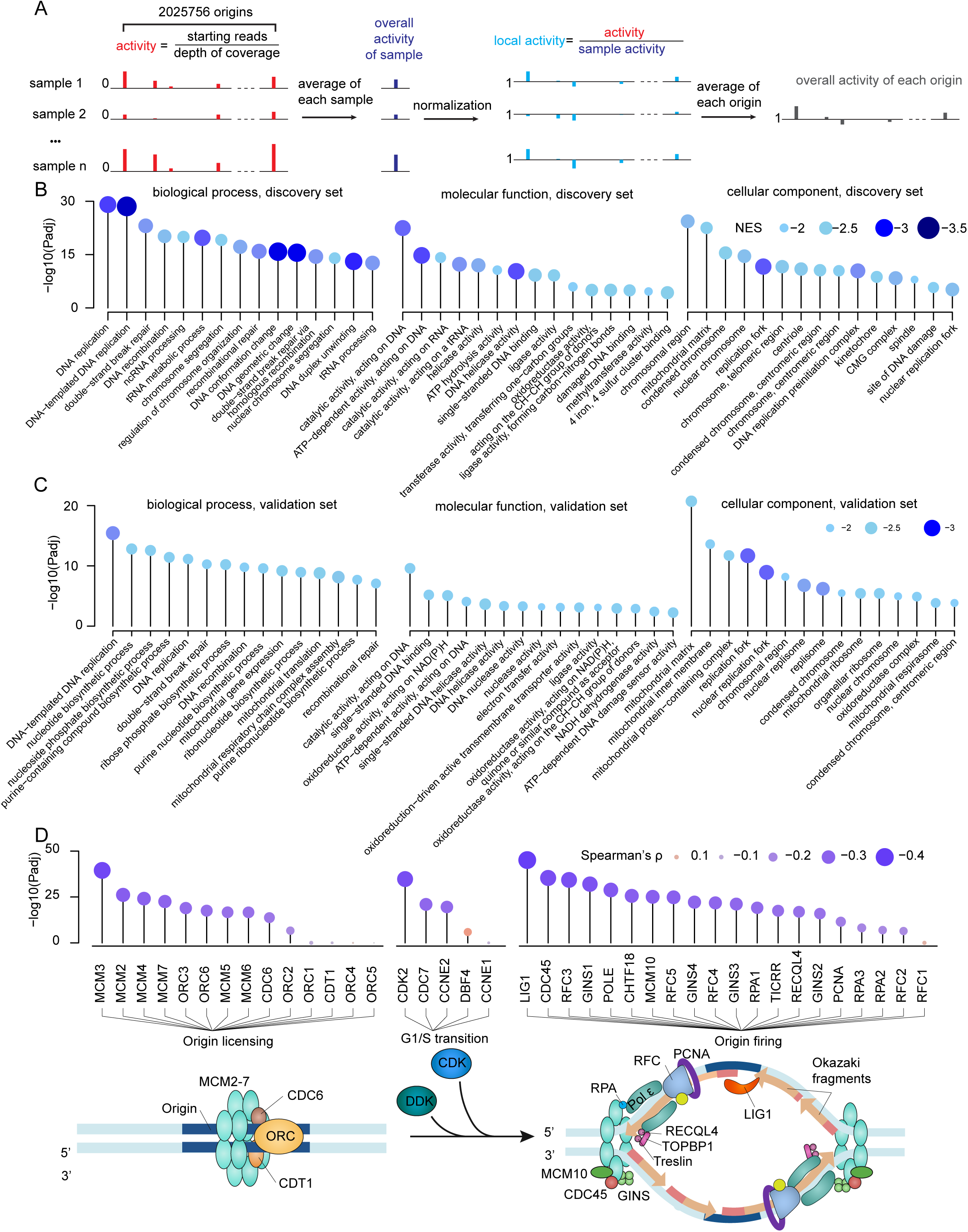
Association of the activity of the origins with gene expression. (**A**) Flowchart of the calculation of the activity of sample and origin. (**B**)-(**C**) The top gene ontology terms associated with the average DNA replication activity per sample by gene set enrichment analysis, on the discovery (**B**) and validation (**C**) datasets. (**D**) (Top) Association of key genes involved in origin licensing, G1/S transition, and origin firing with activity. (Bottom) Model of the process of DNA replication origin licensing to firing. Created in BioRender. Li, Z. (2025) https://BioRender.com/uiw1vxg.

Overall, replication activity was highly variable across the samples (**Fig. S3B**). To identify what was driving this difference, we used the RNA-Seq-based gene expression data of 1308 matched T-ALL samples and performed gene set enrichment analysis (GSEA) of gene ontology terms (*18*). For each gene, we calculated Spearman’s correlation coefficient of the expression of the gene with the average activity of the DNA replication origins, which was used as the ranking metric in GSEA analysis. In this analysis, the biological processes of DNA replication, DNA conformation change and DNA geometric change, the molecular functions catalytic activity acting on DNA and DNA helicase activity, and the cellular components replication fork and DNA replication preinitiation complex exhibited the lowest negative enrichment scores (**Fig. 3B**), indicating these terms were negatively associated with origin activity. To validate these findings, we obtained RNA-Seq data of 227 B-ALL patients in the validation dataset, and similar associations were also seen in this validation dataset (**Fig. 3C**).

We then checked the association between the replication activity and specific genes involved in origin licensing, G1 to S transition and origin firing. Most of the genes demonstrated significant negative correlation with the activity of the origins (**Fig. 3D**). Specifically, all the genes in MCM2-7 complex (critical for both origin licensing and firing), CDK2 (important for G1/S transition), and LIG1 (responsible for the ligation of the leading strand and the last Okazaki fragment) were highly correlated with the activity negatively. Similar associations were observed in the validation set (**Fig. S4**), albeit with reduced significance, likely due to the smaller sample size. These data suggest that the detection of starting/ending reads at the DNA replication origins is driven by slower progression of DNA replication or stalled replication forks, allowing accumulation of newly synthesized DNA across multiple cells, making them detectable by our method. More importantly, the direct association of the activity of the origins with the expression of DNA replication-related genes validates their functional relevance.

### Genome-wide variation of replication activity is linked to DNA replication timing

To understand the genome-wide variation in replication activity, we obtained human DNA replication timing data from a previously published high-resolution Repli-Seq study (*11*), which profiled newly synthesized DNA in 16 sorted S-phase fractions for human embryonic stem cell lines (H1, H9) and colon carcinoma line HCT116. For each of the identified replication origin, we estimated its average activity across 2,616 samples in the discovery cohort after normalization. (**Fig. 3A** and **Methods**). Notably, we observed high replication activity for origins located in early-replicating regions, and low activity in late-replicating regions (**Fig. 4A** and **Fig. S5A-B**). Across the genome, activity of DNA replication origins is positively correlated with the amount of newly synthesized DNA in early S phases at the same region, and negatively associated with late S phases as determined by Repli-seq (**Fig. 4B** and **Fig. S5C-D**). Overall, genome-wide variation of replication activity tracked with replication timing (**Fig. 4C** and **4D**).

**Fig. 4.**
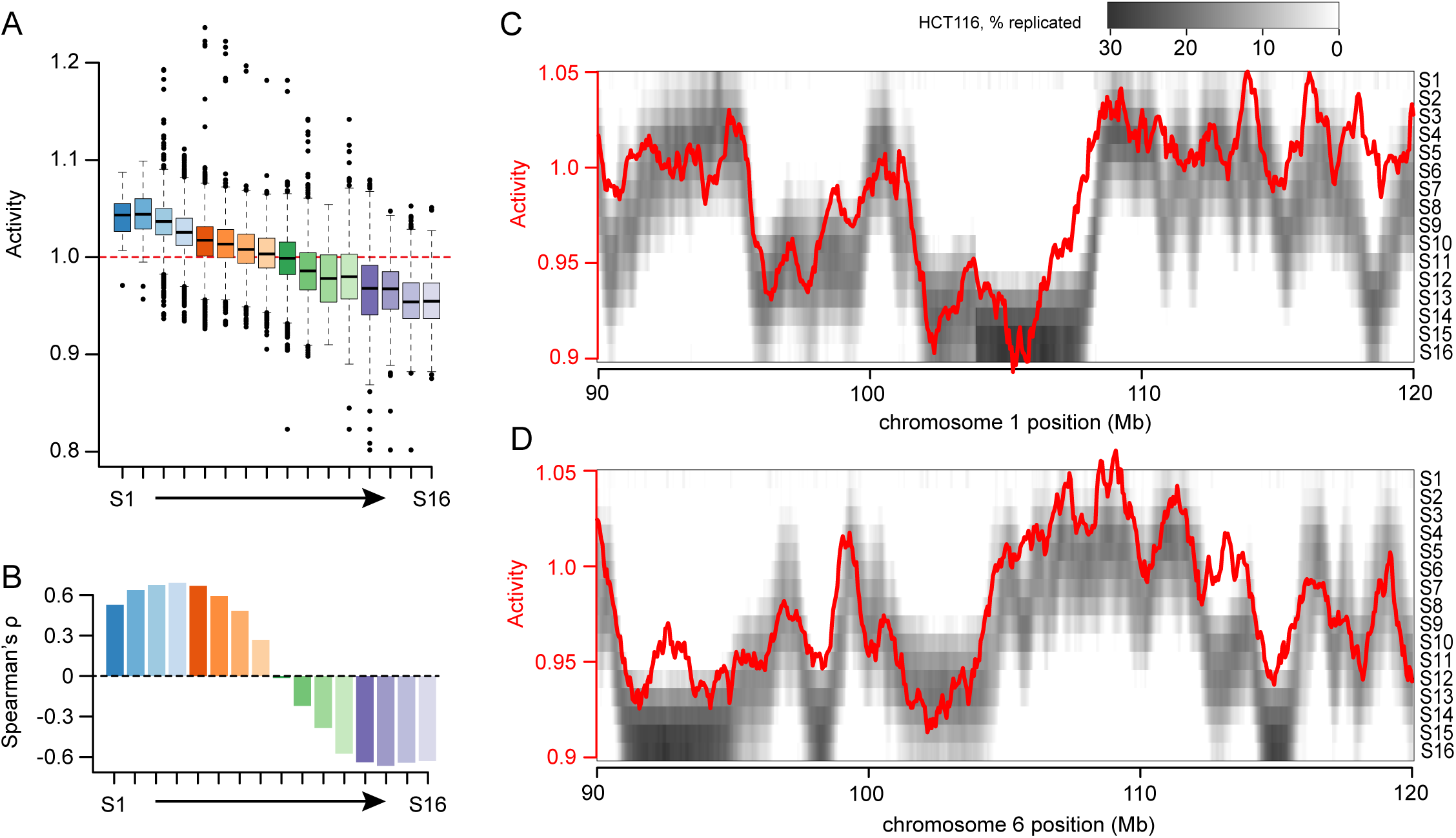
Genome-wide variation of the activity of replication origins. (**A**) Distribution of the average activity of the origins in genomic regions that were replicated during varied S phases based on Repli-Seq data of HCT116 cells. (**B**) Correlation of the average replication activity with the percentage of DNA replicated during varied S Phases in HCT116 cells. (**C**)-(**D**) Two examples showing the concordance between the activity of the origins and Repli-Seq signal of HCT116 cells on chromosome 1 (**C**) and chromosome 6 (**D**).

### Usage of nucleotide alters the activity of DNA replication origins

As demonstrated in **Fig. 1H**, there was considerable conservation in the sequence motif around the replication origins. To further explore this sequence specificity, we comprehensively identified single nucleotide polymorphisms (SNPs) in each origin (−8 to +8 position as defined in **Fig. 1H**) and tested their impact on replication activity (see **Methods**). A total of 114,197 variants were observed for 109,523 origins, for which 2,194 exhibited significant association with replication activity *in cis* (Bonferroni corrected P<0.05; **Fig. 5A**). Notably, polymorphisms at ±1 and ±3 positions are mostly likely related to replication activity and exhibited strong preference in base usage, implying that DNA replication machinery requires some level of sequence specificity. As a result, any genetic variation from the preferred bases (e.g. C, G, or A at position +1; **Fig. 1H**) to a less frequently used base (e.g. T at position +1; **Fig. 1H**) led to decrease in replication activity (**Fig. 5B**). This was also confirmed in 703 unrelated samples in the validation dataset (see **Methods**, **Fig. 5C**).

**Fig. 5.**
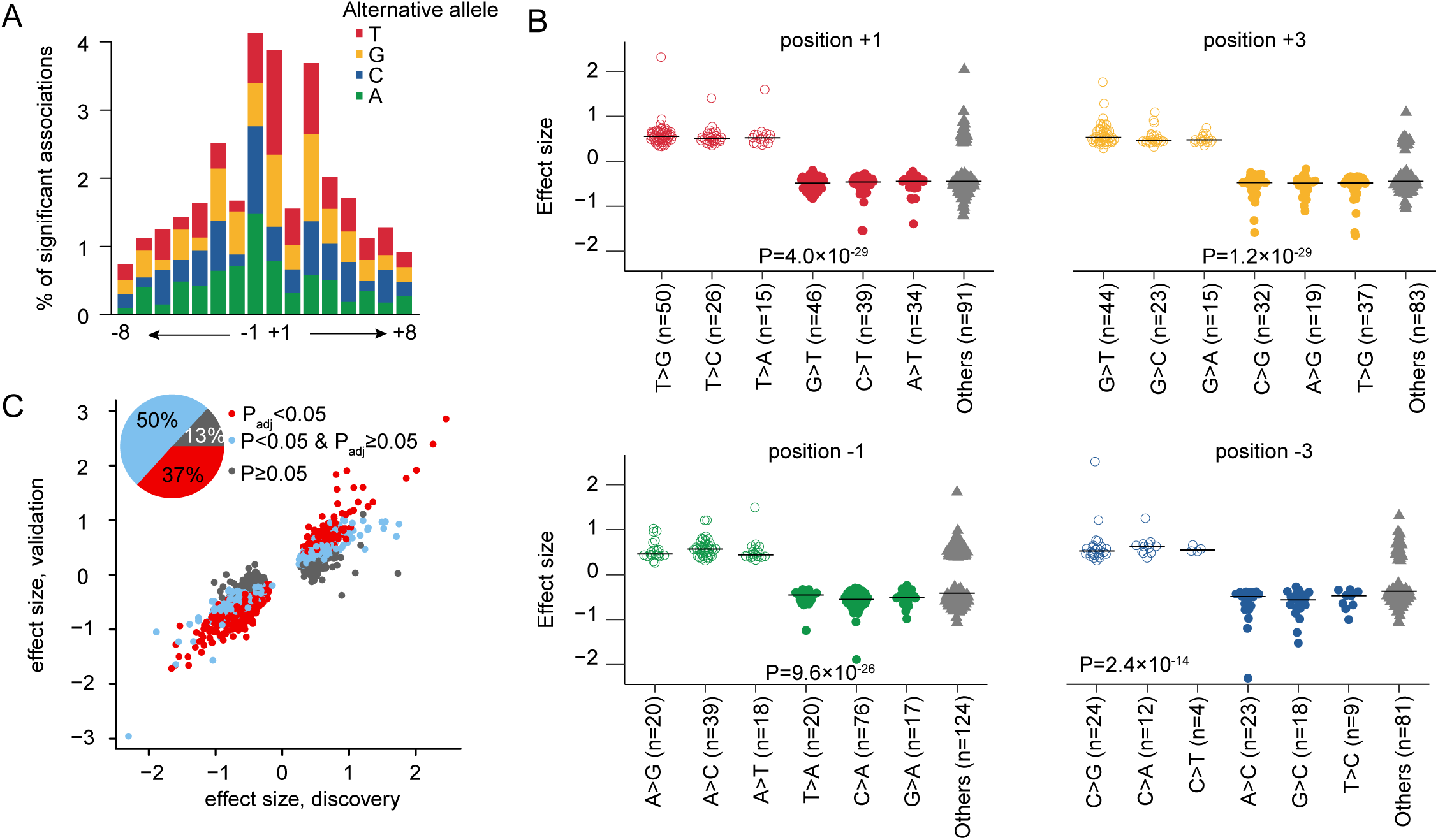
Activity of the replication origins are associated with the nucleotide usage. (**A**) Percentage of SNPs associated with the activity of the origins varied with the relative position of the SNP to the origin. (**B**) In SNPs (within the motif region of the origins) that are associated with the activity of the origins, those with mutation from a rarely used nucleotide in the motif are associated with increased activity while those with mutation to a rarely used nucleotide are associated with decreased activity. (**C**) Validation of the associations on the independent validation set.

### Gene expression is linked to DNA replication activity in cis

Because DNA replication and RNA transcription share the same template, their machinery can interfere with each other, giving rise to correlation between these two types of activities across the genome. To characterize this, we examined replication activity (as 50 kb blocks across the genome) and its association with gene expression (both *in cis* and *trans*), using 1308 T-ALL samples with both WGS and RNA-Seq data. A consistent pattern of *in cis* correlation is identified (**Fig. 6A**, the diagonal line), suggesting that the replication activities of the origins are associated with expression of adjacent genes. In addition, most *in cis* correlations are positive (**Fig. 6A**), i.e., higher gene expression levels are associated with higher activity of the adjacent origins.

**Fig. 6.**
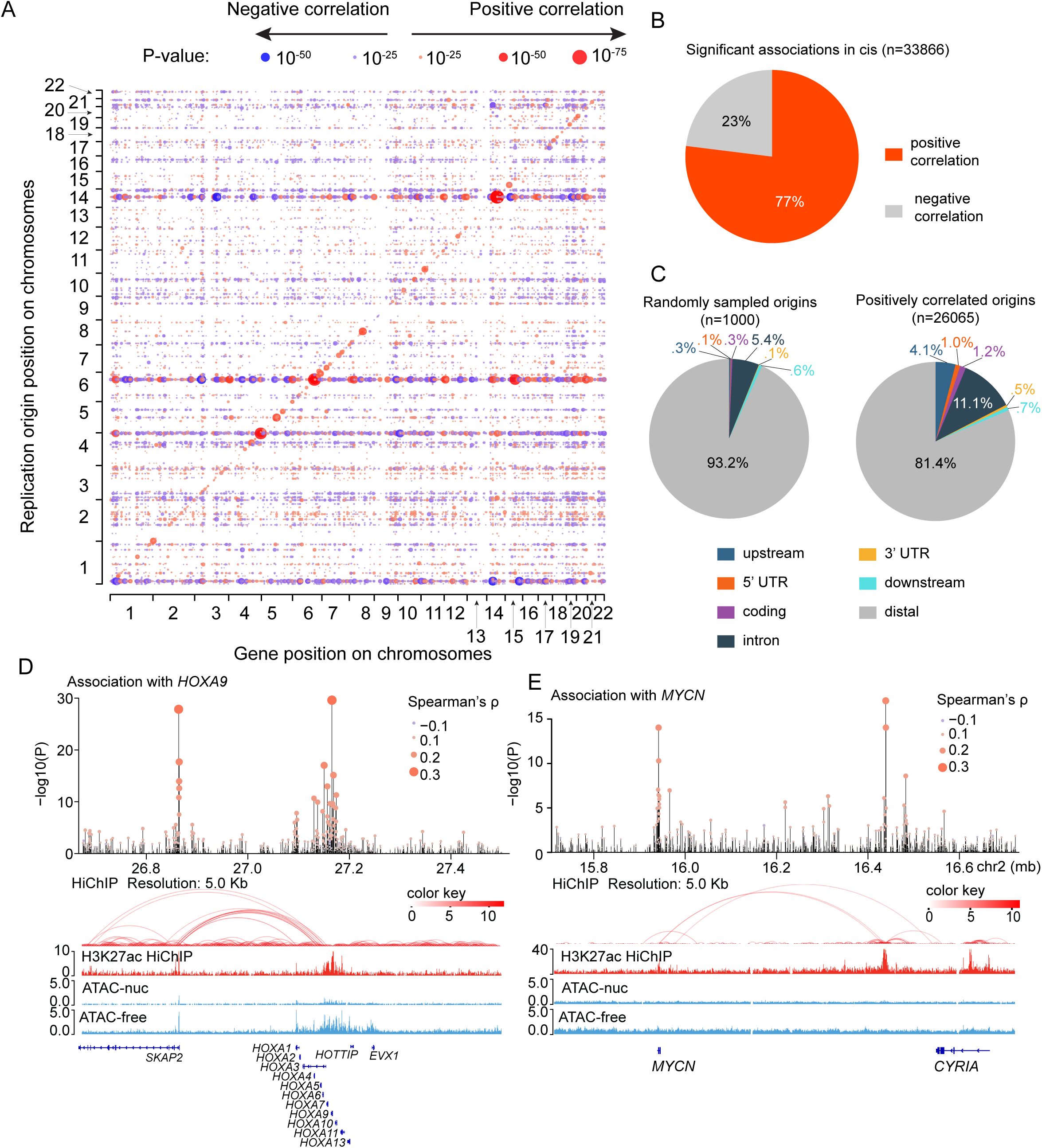
*In cis* association between replication and transcription. (**A**) Significance of the correlation between replication activity per 50 kb window and gene expression revealed *in cis* positive correlation. (**B**) Significant associations between the expression of genes and activity of individual replication origins within 500 kb of the transcription start site of the gene are mostly positive. (**C**) Position of the significantly associated origins relative to the gene. (**D**)-(**E**) Distal association of DNA replication origins with expression of *HOXA9* (**D**) and *MYCN* (**E**) are identified in regions that interact with the respective gene loci.

Focusing on *in cis* correlations, we specifically tested the effect of each replication origin on gene transcription (within ±500 kb of TSS). Of 15,617,355 gene-replication origin pairs tested, 33,866 showed significant associations after Bonferroni correction (P<3.2×10^-9^) with most being positive (77%; **Fig. 6B**). When compared to randomly sampled gene-replication origin pairs tested, replication origins positively associated with transcription are enriched in regions immediately upstream to the gene body (**Fig. 6C**), indicating that transcription factor binding may affect the replication activity in the same region.

This analysis also revealed a large number of distal associations – which may indicate the presence of cis regulatory elements (CREs) linked to the associated gene. For example, at the *HOXA* gene cluster (**Fig. S6**), we identified two genomic loci with multiple origins linked to *HOXA* gene transcription *in cis*, and there was strong evidence for their direct interaction at the 3D chromatin level on the basis of previously published HiChIP data (*12*) (**Fig. 6D**). Similar chromatin interactions involving replication hotspots were also observed at the *MYCN* locus, again supported by HiChIP data (*12*) (**Fig. 6E**).

### Distinct genetic variation patterns around replication origins

To investigate if genetic variation in regions with DNA replication origins are associated with human traits, we examined variants documented in the GWAS (Genome-Wide Association Study) catalogue (*19*). As shown in **Fig. 7A**, there was a general under-representation of genetic polymorphisms proximal to DNA replication origins. By contrast, these regions appear to be enriched for trait-associated variants (**Fig. 7B**). To further explore the impact of DNA replication origin on mutagenesis process in the genome, we analyzed 671,646 *de novo* mutations from the Gene4Denovo database (see **Methods**) (*20*). An increased mutation rate was observed around DNA replication origins (**Fig. 7C**), consistent with a previous report (*21*). Within the sequence motif of the replication origins, the frequency of common SNPs, traits-associated SNPs and *de novo* mutations fluctuated greatly, yet with a consistent pattern, with increased frequency of mutations at positions ±1, and decreased frequency of mutations at positions ±4 (**Fig. 7D**).

**Fig. 7.**
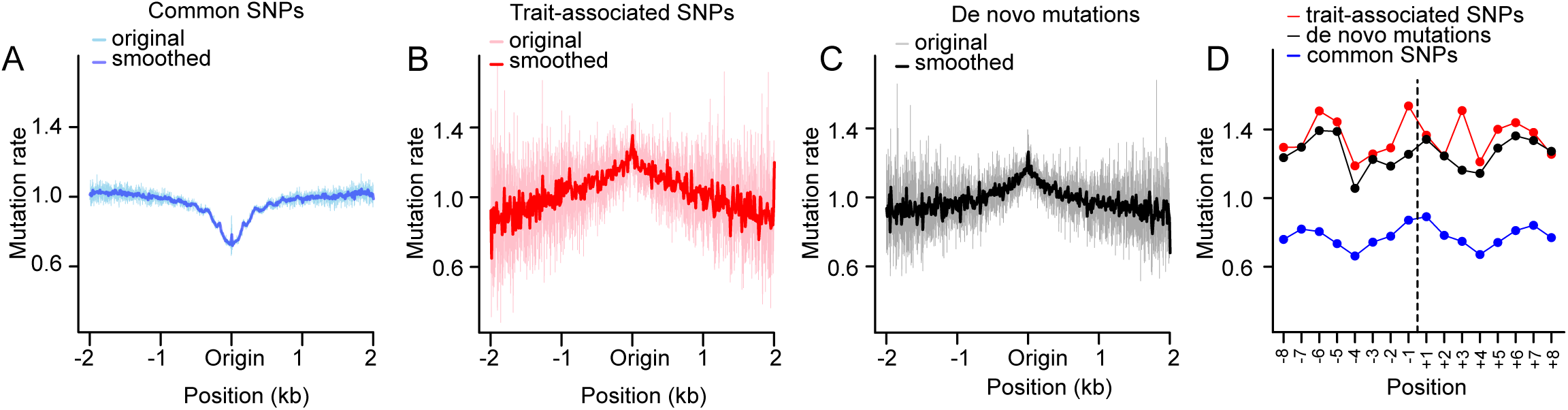
DNA replication origin is linked to phenotypic heterogeneity and *de novo* mutations. (**A**) Frequency of common SNPs is lower around DNA replication origins. (**B**) Frequency of trait-associated SNPs is increased in regions adjacent to DNA replication origins. (**C**) Frequency of *de novo* mutations is higher around DNA replication origins. (**D**) Variations of the frequencies of common SNPs, trait-associated SNPs and *de novo* mutations in the 16 bp motif region of DNA replication origins.

## Discussion

It has long been thought that DNA replication origins of metazoan genomes are defined by large scale sequence and structural characteristics and lack a specific motif. Here, using commonly available WGS data, we describe the exact site of DNA replication origins in the human genome, which are mapped to more than two million genomic positions. Our method of replication origin identification was based on the observation that SRE and SRS events preferably occur next to each other. This observation strongly suggested that the two leading strands start at the exact same position, similar to what is seen in the autonomously replicating sequence (ARS) 1 of budding yeast (*22*). We further showed that the DNA replication origins exhibit a 16 bp motif, which is symmetric, palindromic, and reverse-complementary. While this motif is likely related to the specificity of the protein-DNA binding at DNA replication origins, the exact mechanism that dictates this sequence usage requires further study.

The origins identified by our methods are supported by multiple types of evidence. First, the density of the origins varies consistently with replication origins identified by previous methods such as SNS-Seq and Ini-Seq2. Second, the activity of the origins by WGS is associated with the expression of DNA replication-related genes. Lastly, the genome-wide variation of the replication activity are correlated with replication timing. Collectively, our data suggest that the origins we identified probably reflect stalled or slowly progressing replication forks and are enriched in early replicating regions. This can be linked to low expression of DNA replication-related genes, slow DNA synthesis during early S phase, resulting in the accumulation of newly synthesized DNA (across multiple cells) at early origins.

It should be noted that the ∼2 million DNA replication origins identified in this report constitute only a subset of the origins in the human genome. The 16 bp motif of DNA replication origins is highly degenerate with a normalized Shannon entropy of 0.81. Occurrence of the motif sequence is ubiquitous in the genome, which may be necessary for the rapid replication of the genome. Due to the low complexity of the sequence motif, it is understandable that previous DNA replication origin mapping methods, mostly based on read density, are not able to identify the exact position of DNA replication origin and the sequence motif. On the other hand, CHiP-Seq of ORC or double hexamer MCM may be able to identify the most consistently licensed origins, but they are not necessarily those that are consistently fired. In S phase, MCM and other components of the DNA replication machinery can essentially bind anywhere in the genome. Instead of relying on read density, our method uses read sequence (especially at the start and/or end of reads), thereby providing a new way to identify the exact starting position of DNA replication.

Because DNA replication is a fundamental biological process, identifying the exact sites of replication origin is of immediate importance. DNA replication is closely linked to gene expression (*23*), DNA repair (*24*), *de novo* mutations (*21*), as well as genomic alterations in cancer (*25, 26*). Our data imply potential impact of DNA replication origins on human traits and/or local mutation rate, although the mechanisms underlying this relationship remain to be determined. Because our method is dependent on sequencing of non-ligated newly synthesized leading strands, it is expected that the chemistry used in sample processing, DNA extraction, WGS library construction and next generation sequencing would affect the detectability of the origins. As the newly synthesized DNA are shorter, it is important that small DNA are retained in the process. It can also be postulated that unbiased DNA fragmentation methods, such as sonication rather than enzymic fragmentation, should be used. Meanwhile, all samples analyzed in this study were derived from cryopreserved cells, which likely helped maintain their cellular state prior to DNA extraction. Our method can also be easily applied to other species. To show that, we applied our method to the WGS data of the spleen samples of three mice, and a similar motif was identified (**Fig. S7**).

In conclusion, we identified the exact sites of DNA replication origins in the human genome, which exhibit a 16 bp consensus motif. Activities of these origins are associated with DNA replication efficiency and timing. These results provide important insights into fundamental processes in cell biology and genomics.

## Methods

### Data and preprocessing

The primary analysis of this study was based on a published dataset of WGS and RNA-Seq data of pediatric patients with T-ALL (*12*). This dataset includes WGS data of 1308 leukemic samples collected at diagnosis and 1308 normal samples collected at remission, of which 1307 samples are paired. RNA-Seq data are also obtained for the 1308 leukemic samples. A validation dataset was obtained from Gabriella Miller Kids First Pediatric Research Program Down Syndrome (DS), Heart Defects, and Acute Lymphoblastic Leukemia study. This dataset consists of a total of 1182 samples, including 1) WGS data of 249 samples of DS children with congenital heart disease, and 120 samples from their parents; 2) WGS data of 359 leukemic samples and 454 normal samples, as well as RNA-Seq data of 227 leukemic samples of DS children with B-cell ALL. Other data used in this study for different analysis performed are reported in the following Methods sections.

For WGS data, reads were mapped to hg38 genome reference using bwa (version 0.7.15-r1140) (*27*). Joint variant calling was performed using GATK (version 4.0.10.1) (*28*). For RNA-Seq, reads were mapped to hg38 genome reference using STAR (version 2.7.1a) (*29*) and the number of reads mapped to each gene were counted using RSEM (version v1.3.3) (*30*). Gene expression levels were then normalized using variance stabilizing transformation from DESeq2 package (version 1.44.0) (*31*).

#### Identification of the exact sites of DNA replication origins from WGS data

In WGS data, the number of reads starting (denoted as *S*) or ending (denoted as *E*) at a genomic position follows a binomial distribution with the following probability mass function:

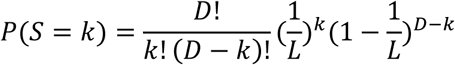

Here, *L* is the length of the reads and *D* is the depth of coverage at the position.

Given the WGS data of a sample, we first scanned the genome to record the total number of reads starting at (*s*_*i*_), ending at (*e*_*i*_), and covering (*d*_*i*_) any given position *i*. We define a position as SRS event when

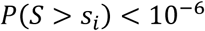

and as SRE when

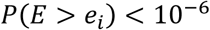

An SRE-SRS pair is defined for position *i* when

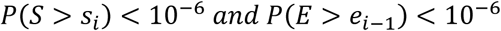

As a first cut, we require *s*_*i*_ and *e*_*i*−1_ to be at least four reads. Only genomic regions not in repeatmasker or ENCODE blacklist (*32*), and with 100 bp mappability (*33*) of 100% were included in the analysis. Only autosomes are included in the analysis. Overall, a total of 1,296,046,051 bp of the genome was analyzed.

To show that these SRE-SRS pairs were not the biased breakpoints during fragmentation, we analyzed the relative depth of coverage at these positions. The depth of coverage of all the identified DNA replication origins were extracted, which were divided by the average depth of coverage of the sample, estimated from sampling 10,000 random positions in the analyzed region mentioned above.

### Activity of DNA replication origins

To measure the detectable activity of DNA replication origins, we counted the number of reads starting at position +1, this number was then divided by the total number of reads covering this position. Hence, for the *i*th origin in the *j*th sample, the raw activity was calculated as follows:

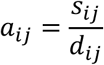

Similarly, ending reads-based activity was calculated by the number of reads ending at position - 1 divided by the total number of reads covering position −1. For each sample, the average activity calculated over all the origins as:

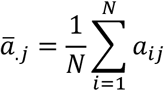

where *N* = 2,025,756 is the total number of origins. To capture the genome-wide variation of DNA replication origin activities, we normalized the raw activity by dividing the average for each sample:

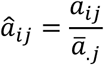

Then for each origin, we can obtain the averaged normalized activity as follows:

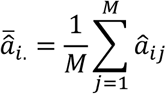

where M is the number of samples included in the analysis. We consider *a̅*_.*j*_ as the overall activity of sample *j*, and 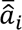 as the activity of origin *i*.

### Gene expression analysis

The correlation between gene expression and average activity of the samples (*a̅*_.*j*_ as described above) were assessed using Spearman’s correlation coefficient and statistical significance of the correlation was tested using Student’s t-test. GSEA of gene ontology terms was performed using the gseGO function from clusterProfiler (version 4.12.6) (*34*) package, using correlation coefficient as the ranking criteria. For both discovery and validation datasets, this was performed on samples for which both WGS and RNA-Seq data were available. Specifically, for the primary discovery set, 1308 T-ALL diagnostic samples were included; for the validation set, 227 DS-ALL diagnostic samples were analyzed.

Association between local replication activity (normalized activity *â*_*ij*_ as described above) and gene expression were performed similarly using correlation coefficients. To reduce the number of tests, we first assessed the correlation between gene expression of all genes with the average activity of DNA replication origins within each 50 kb window, which revealed a pattern of *in cis* association. We then analyzed the correlation of each gene with the activity of all identified DNA replication origins with 500 kb of its transcription start site.

### Replication timing

Human replication timing data was obtained from a previous publication by Zhao et al (*11*). This study sequenced the amount of newly synthesized DNA in 16 equally partitioned S phases, namely S1 to S16 (early to late). The Repli-Seq heatmap matrixes of human cell lines H1, H9 and HCT116 were downloaded from gene expression omnibus (id: GSE137764), and values of each 50 kb bin are scaled linearly to have a total of 100, so that the values reflect the percentage of DNA being synthesized in a specific S Phase.

For each 50 kb bin across the genome, we assigned the genomic region to one of the 16 S phases with the highest value. We calculated the average of the normalized activities (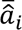. above) of DNA replication origins in each 50 kb bin. Only bins with at least 10 detected origins were included in this analysis. The activity of each 50 kb bin is then smoothed by taking the average of the neighboring bins with distance less than 500 kb, which is used to compare with replication timing data.

### Association between single nucleotide polymorphisms and DNA replication origin activity

For each DNA replication origin, we evaluated the association between normalized activity (*â*_*ij*_ as described above) and single nucleotide polymorphisms present within its ±8 bp motif region. Linear regression was performed with the normalized activity as dependent variable, and the genotype (number of alternative alleles) as independent variable, controlling for age, sex, as well as the first 10 principal components of the genotype data to account for genetic ancestry. Principal component analysis of the genotypes was performed using plink2 (version 2.3) (*35*). Age is not included in the validation analysis due to unavailability of age information. Only binary SNPs with high genotyping rate (>99.9%) and allele frequency larger than 1% were included in the analysis. For primary discovery data, this analysis was performed only on the normal samples (n=1308). For validation dataset, this analysis was performed on a total of 703 samples, including the normal samples of DS-ALL patients (n=454), and samples of DS-CHD patients (n=249). Variant on chromosome 21 were not analyzed for validation as most of the samples have trisomy 21. The p-values were adjusted for multiple testing using Bonferroni correction with adjusted P<0.05 considered as statistically significant. SNPs that demonstrated significant association with the activity of the origins in their neighborhood were then tested on the validation cohort.

### De novo mutation and trait-associated variant analysis

*De novo* mutations were downloaded from the Gene4Denovo database (*20*), and trait-associated variants were obtained from the NHGRI-EBI Catalog of human genome-wide association studies (*19*). Common SNPs (with allele frequency >1%) obtained from dbSNP build 156 were analyzed for comparison (*36*). Only mutations/variants on autosomes were included. For *de novo* mutations, because this data set is based on hg19, the genomic positions of replication start sites were lifted to hg19 from hg38 using loftOver tool, and only those with a reference allele same as the genome reference were included (total 671,646 mutations). For trait-associated variant analysis, only SNPs achieved genome-wide significance (P<5×10^-8^; total 295,643 SNPs) were included in the analysis. A total of 1,7181,810 common SNPs were analyzed.

For genomic positions that are *i* bp apart from the DNA replication origins (that is, the distance to the closest DNA replication start site is *i* bp), we calculated the enrichment score of mutations using the following formula:

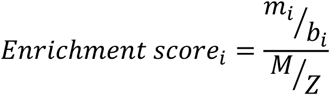

In this formula, *m*_*i*_ is the number of mutations detected at these positions, *b*_*i*_ is the number of genomic positions with a closest distance of *i* bp to DNA replication start sites, *M* is the total number of mutations and *Z* is the ungapped genome size (2,756,349,822 for hg38 and 2,684,578,480 for hg19).

## Acknowledgements

We thank Dr. Angela McArthur from the Department of Scientific Editing at St. Jude for proofreading. **Funding:** This work was in part supported by the National Institutes of Health (X01HL145686-01, X01HD100702, R01CA249867, U01HD116485, P30CA125123-14S4, 1R03HD103908-01 and R03CA256550), Department of Defense (W81XWH-20-1-0567), the Lynch family, and the American Lebanese Syrian Associated Charities. **Author contributions**: Study concept and design: Z.L. and J.J.Y. Acquisition of data: S.Y., C.G.M., T.G.R., E.J.L.-C., S.L.S., K.R.R., P.J.L., D.T.T. and J.J.Y. Data analysis and interpretation: Z.L., W.Y., G.W., and J.J.Y. Drafting of manuscript: Z.L. and J.J.Y. Reviewing and editing of manuscript: All authors. **Competing interests**: The authors declare no competing interests. **Data and material availability**: The sequencing data (whole-genome and RNA-Sequencing) of both the primary discovery and the validation cohorts are available from the database of Genotypes and Phenotypes (dbGaP) under accession numbers phs002276.v2.p1 and phs002330.v2.p1, respectively.

## List of Supplementary Materials

Supplementary Materials

Fig. S1 to S7

