## Supplemental figures for "The Genomic Architecture of Human DNA Replication Origins"

**Li et al.**

Supplementary figures

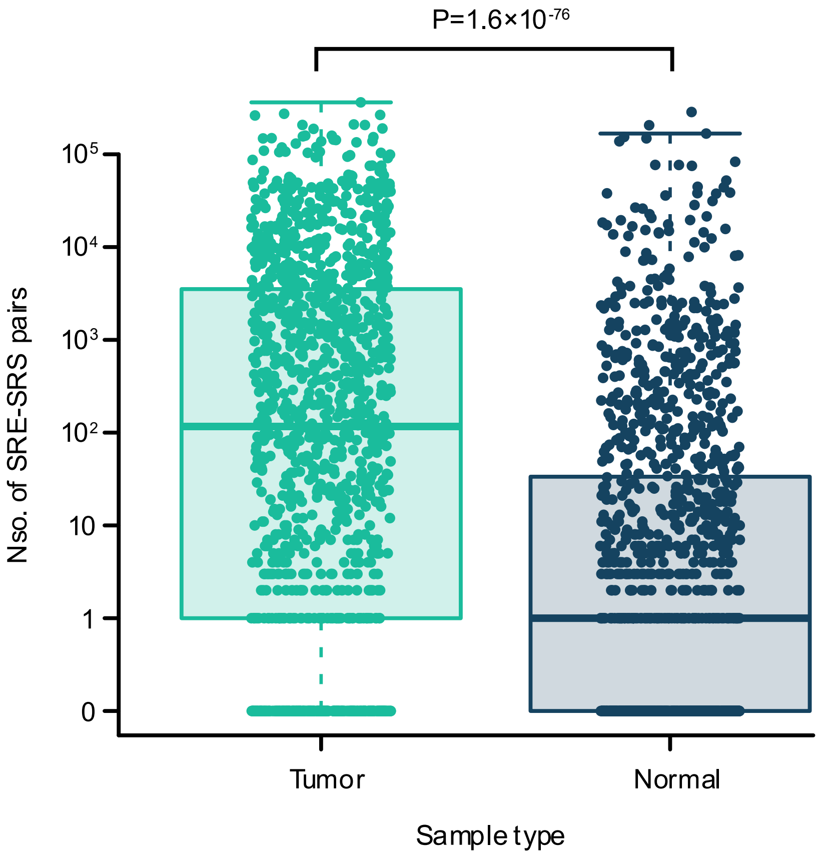


Fig. S1. Comparison of the number of SRE-SRS pairs detected in tumor and normal samples of T-ALL patients. P-value is calculated with unpaired Mann–Whitney U test.

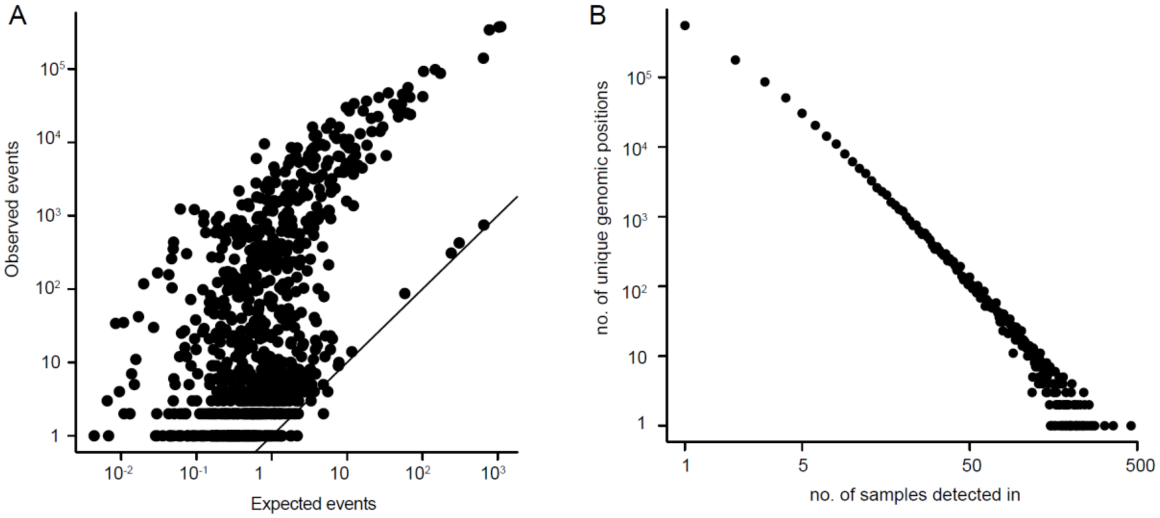


Fig. S2. Identification of DNA replication origins on the validation data. (A) number of identified SRE-SRS pairs are higher than expected. (B) distribution of the number of samples an origin is detected in follows an exponential distribution.

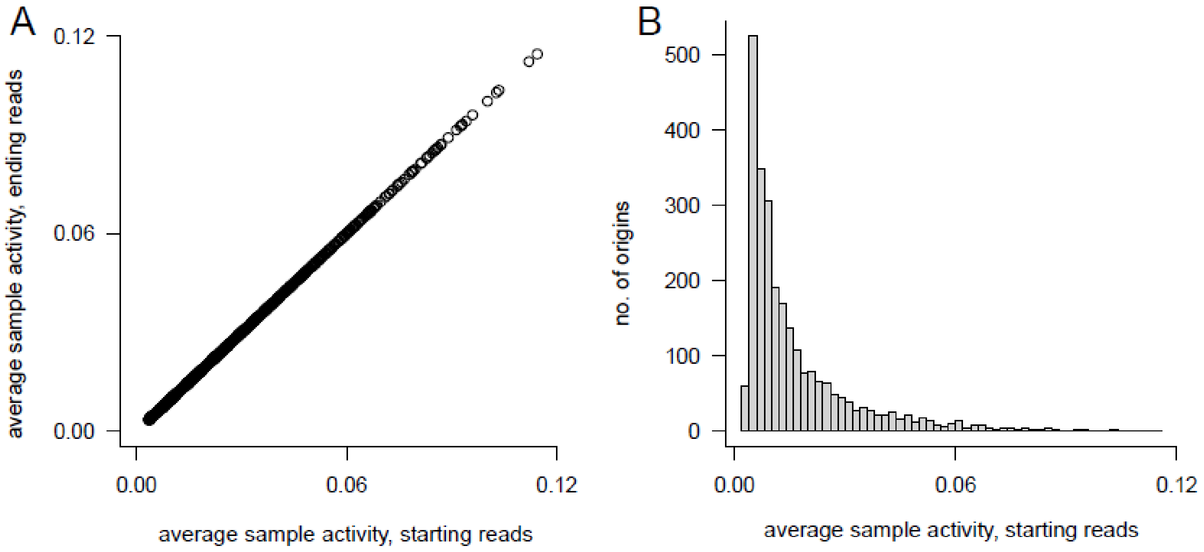


Fig. S3. Average detected activities of the samples. (A) starting reads-based and ending reads-based activities are highly correlated. (B) distribution of the average activities of the samples.

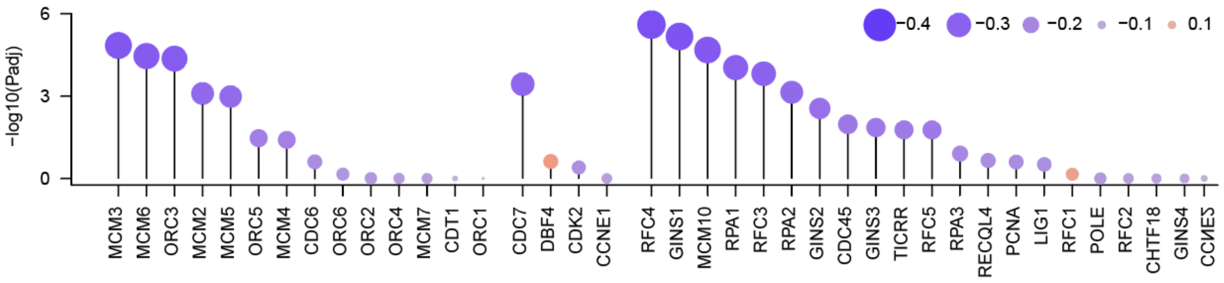


Fig. S4. Association of the DNA replication origin licensing, G1/S transition and origin firing genes with the average activity of DNA replication origins on the validation data.

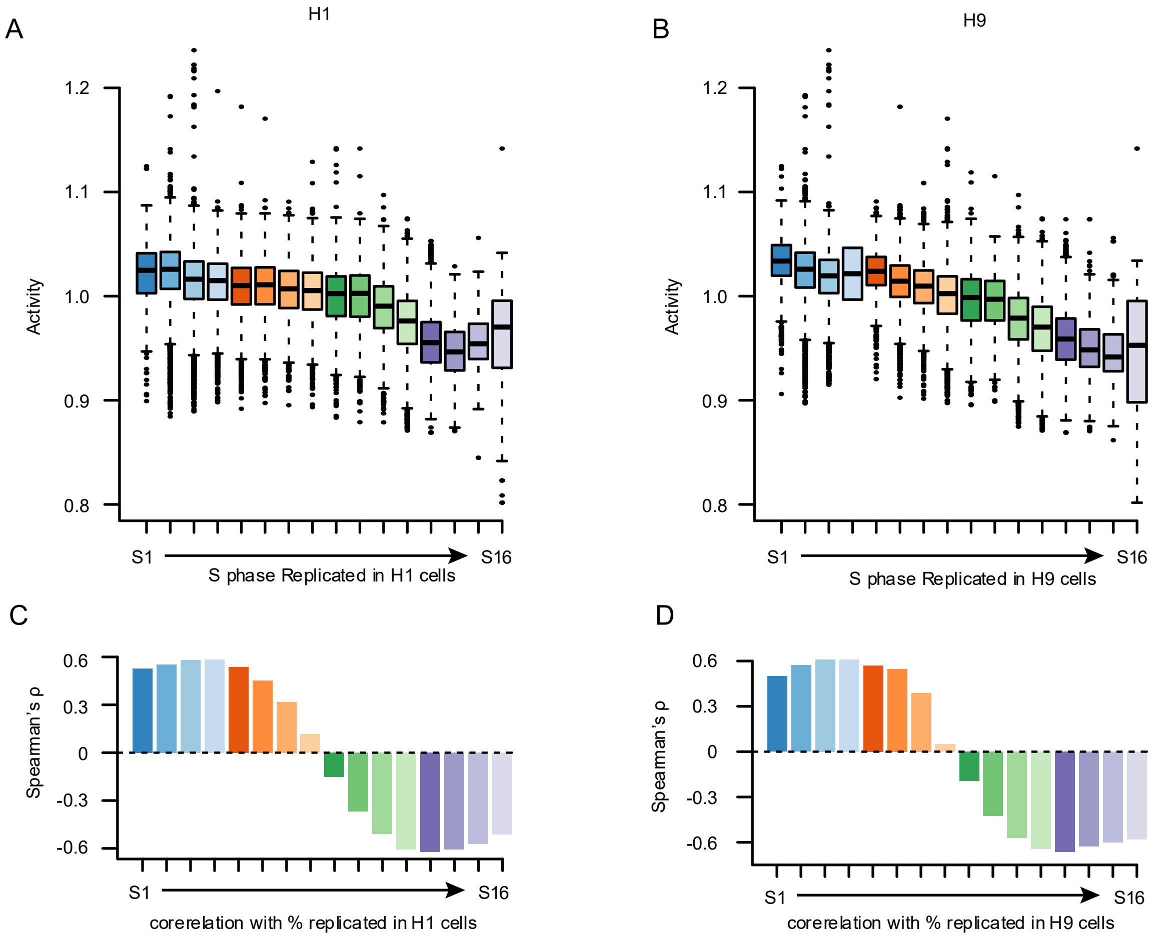


Fig. S5. Association the activity of replication origins with replication timing in H1 and H9 cells. A.-B. Distribution of the average activity of the origins in genomic regions that were replicated during varied S phases based on Repli-Seq data of (A) H1 and (B) H9 cells. C.-D. Correlation of the average replication activity with the percentage of DNA replicated during varied S Phases in (C) H1 and (D) H9 cells.

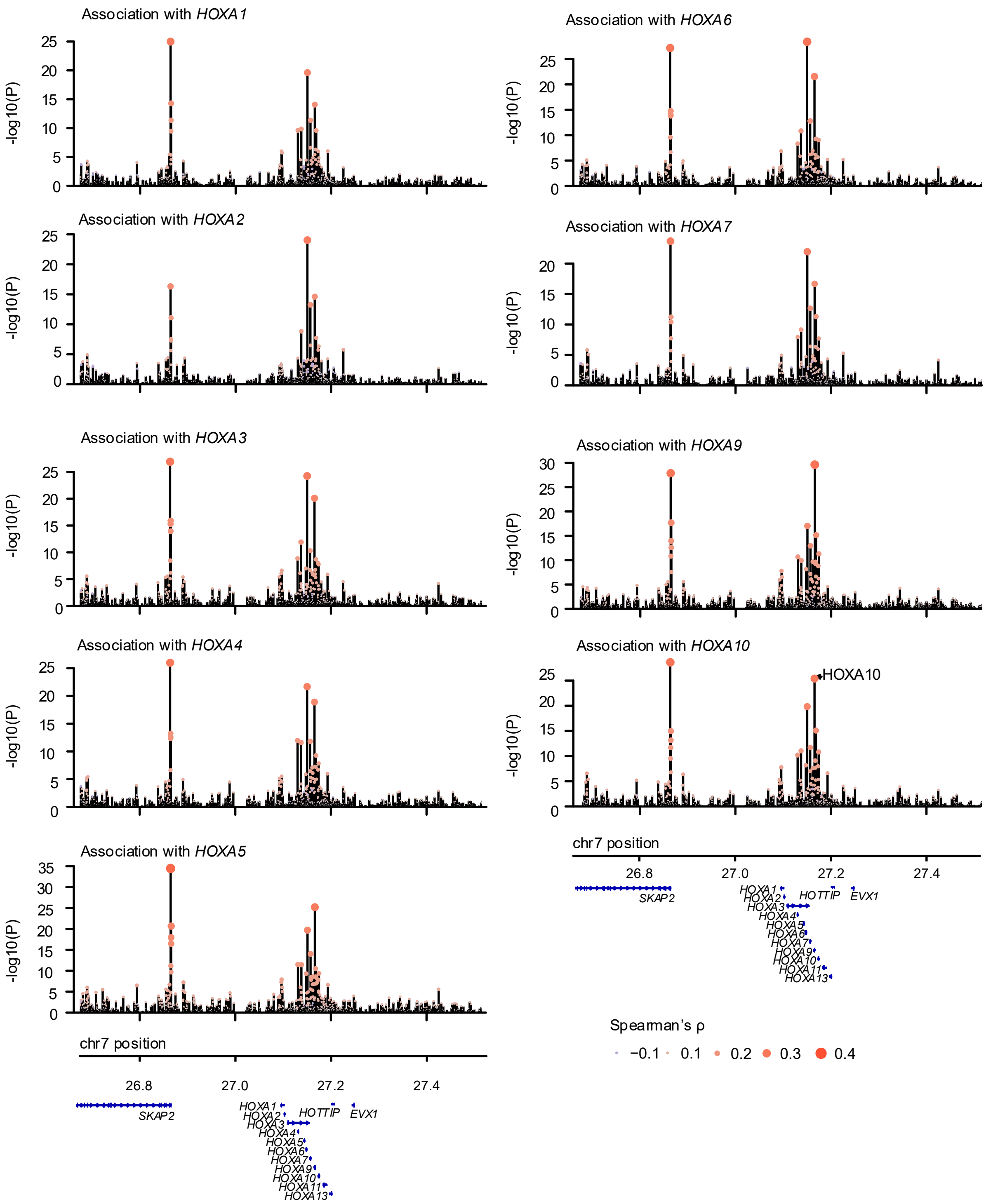


Fig. S6. *In cis* association of the activity of DNA replication origins with the expression of HOXA genes.

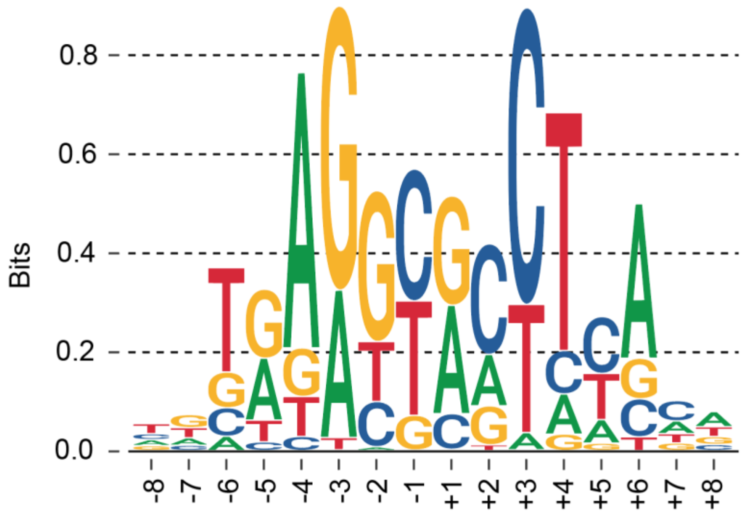


Fig. S7. The motif of DNA replication origins identified in mice. Whole genome sequencing was performed on spleen samples of three mice. Adjacent ending and starting events were identified using the same methods as described in Methods, with the only exception that a higher threshold of P-value (0.001) was used for significant reads starting (SRS) and significant reads ending (SRE) events identification.
